# Cell Type Weighted Dimensionality Reduction

**DOI:** 10.64898/2026.04.30.721796

**Authors:** Santosh Putta, Wesley Jensen, Srikar Devakoda, Leesa Pennell, Josh Croteau

**Affiliations:** Revvity Inc.

**Keywords:** Single-cell, dimensionality reduction, cell-lineage, visualization, data analysis, non-linear projections

## Abstract

High-dimensional single-cell technologies, such as flow cytometry and CITE-Seq, typically rely on established lineage markers to define cell identities. Additional markers are commonly analyzed within the context of these predefined cell types. Nonlinear projection methods such as t-SNE and UMAP provide a visual framework for this analysis by enabling the overlay of cell types and marker expression. However, these methods frequently produce projections where distinct cell types substantially overlap, hindering interpretation of marker expression patterns relative to known cell types.

In this study, we investigate the underlying causes of this phenomenon and demonstrate that such overlaps often stem from the inherent high-dimensional structure of the data rather than limitations in the dimensionality reduction algorithms themselves. To address this, we introduce Cell Type Weighted Dimensionality Reduction (CWDR), a novel approach that incorporates lineage-based information through a supervised weighting mechanism.

By integrating both cell identity and marker expression, CWDR preserves the visual separation between predefined cell types while maintaining the local variance necessary for downstream analysis. We validate our method across multiple high-dimensional flow cytometry and proteogenomic datasets. Our results show that CWDR significantly reduces inter-cluster overlap compared to traditional methods, providing a clearer framework for visualizing marker expression within the context of specific cell lineages.

## 1 Introduction

Advances in single-cell technologies such as spectral flow cytometry and CITE-Seq now enable the quantification of tens to hundreds of proteins and thousands of RNA transcripts across thousands to millions of individual cells. When analyzing samples from healthy individuals, disease cohorts, or experimental model systems, investigators typically begin by leveraging established knowledge of cell-lineage and function. As a result, major immune cell populations—such as T cells, B cells, and their respective subtypes—are routinely identified using canonical protein markers; for example, T cells are defined by expression of CD3. Domain experts commonly assign cell identities through hierarchical gating, in which protein expression patterns are examined in two-dimensional plots. This classification provides essential biological context, as downstream analyses and novel findings are most interpretable when framed within well defined cell types and subtypes. Although cell typing typically relies on a subset of measured markers, it plays a foundational role in organizing high-dimensional single-cell datasets.

It is now possible to measure hundreds to thousands of additional protein and RNA markers in single-cell experiments. Analyzing these markers within the context of established cell types and relevant biological or clinical metadata provides substantial interpretive value. Consequently, it is common practice to quantify protein, RNA, and other markers—particularly those not used for cell-type assignment—within each cell type for downstream analysis. This quantification is often complemented by visual inspection of the single-cell data to assess heterogeneity and co-expression patterns across cell types. However, the high dimensionality of these datasets poses challenges for visualization. Dimensionality reduction methods such as t-SNE and UMAP are therefore routinely applied to project single-cell measurements into lower dimensions for visualization of cellular structure and marker expression. These nonlinear methods aim to learn the underlying high-dimensional manifold and produce a lower dimensional approximation that preserves local relationships between cells, while allowing distortion of long-range distances.

While dimensionality reduction provides a convenient means to visualize high-dimensional data in lower dimensions, it is typically performed independently of cell-type definitions. As a result, cell types frequently overlap in the projection space (e.g., Figure 4a and 4c), complicating the interpretation of marker expression within and across populations. Despite this common observation, the underlying reasons for such cross-phenotype overlap remain insufficiently understood. Here, we investigate the root causes of this phenomenon and propose an enhanced dimensionality reduction framework that explicitly incorporates cell phenotype information.

## 2 Results

We conducted our analyses on high-dimensional single-cell datasets generated using flow cytometry and CITE-Seq. Our objectives were twofold: (1) to determine why lineage-defined cell types frequently overlap in low-dimensional projection space, and (2) to develop an improved dimensionality reduction approach that accounts for biologically meaningful cell types. To achieve these aims, we manually assigned lineage-based cell types, quantified the extent of overlap, and then used these insights to design a more suitable dimensionality reduction framework for single-cell data.

### 2.1 Assigning cell types

The results presented in this manuscript are derived from single-cell protein data generated using flow cytometry and CITE-Seq. The flow cytometry dataset includes 25 protein measurements plus two scatter parameters obtained from 373115 peripheral blood mononuclear cells (PBMCs), while the CITE-Seq dataset comprises 153 protein measurements from 12,456 PBMCs. In both datasets, cell types were assigned through hierarchical gating using cloud software MAS^3^ and CytoScribe^4^ using established lineage markers. A total of 46 and 63 cell types were defined for the CITE-Seq and flow cytometry datasets, respectively, spanning T, B, NK, and monocyte subtypes. The complete list of cell types and gating strategies is provided in the Supplementary Materials (Supplemental Figures A and B).

### 2.2 Investigating overlap of cell types in projection space

We hypothesized that the overlap, or “smudging,” of cell types in projection space may arise from two distinct causes. First, non-linear dimensionality-reduction methods such as t-SNE face inherent challenges when projecting high-dimensional data into low-dimensional space and may therefore only partially preserve neighborhood structure. Second, lineage-defined cell types may already exhibit substantial overlap in their local neighborhoods in the original high-dimensional space, such that inter-cell distances do not sufficiently separate them. In this latter scenario, the dimensionality-reduction algorithm may in fact perform as intended yet still be unable to produce well-separated clusters due to the intrinsic structure of the data. To distinguish between these two possibilities, we analyzed high-dimensional flow cytometry and CITE-Seq datasets.

#### 2.2.1 Quantifying cell type overlap

To quantify cell-type overlap in both the original high-dimensional space and the projection space, we used neighborhood purity (*N*_*P*_), a conventional metric defined for each cell type *C. N*_*P*_ represents the fraction of a cell’s neighbors that also belong to *C*. A cell type which does not overlap with other cell types should have very few neighbors of other cell types, therefore resulting in a *N*_*P*_ value close to 1. However, a cell type with significant overlap with other cell types should have an *N*_*P*_ value closer to zero. The neighboring cells for a cell type are defined as follows:

1. For each cell in cell type *C*, we find the *k* nearest neighbors and enumerate the unique set *S*_*U*_ of these neighbors.
2. Enumerate subset *S*_*UC*_ of *S*_*U*_ that belong to *C*

Then neighborhood purity *N*_*P*_ is the ratio of the number of cells in *S*_*UC*_ and *S*_*U*_ ; i.e.

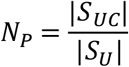

The *k* nearest neighbors for each cell were enumerated using Faiss^5^ library with Euclidean distance metric and *k=20*, consistent with default settings for t-SNE.

Note that *N*_*P*_ can be computed in the original high-dimensional space and in the projection space. Therefore, neighborhood purity values for the cell types in the original high dimensional space can be compared to the corresponding values in the projection space. Since the primary goal for dimensionality reduction algorithms like t-SNE is to preserve the neighborhood, one should expect that the *N*_*P*_ values are similar between the high-dimensional and low-dimensional space if the algorithms performed well. On the other hand, if the *N*_*P*_ values are low in the original high dimensional space, it is unlikely that dimensionality reduction methods will be able to separate the cell types from each in the low dimensional space.

Neighborhood purity (*N*_*P*_) can be computed in both the original high-dimensional space and the corresponding low-dimensional projection. Comparing *N*_*P*_ values across these spaces enables assessment of how well a dimensionality-reduction method preserves local neighborhood structure. Because algorithms such as t-SNE are designed to maintain neighborhood relationships, *N*_*P*_ values in the projection space should closely match those in the original space when the method performs well. Conversely, if a cell type exhibits low *N*_*P*_ in the high-dimensional space, dimensionality-reduction algorithms are unlikely to separate that cell type from others in the projection, regardless of parameter settings.

#### 2.2.2 Distinct cell types often overlap in the original space

We evaluated neighborhood purity for each cell type in the original high-dimensional space. As shown in Figure 2, several cell types exhibit low purity, indicating that their neighborhoods— when computed across all dimensions—contain a mixture of cells from both the target cell type and other types. In contrast, purity values are higher for terminal cell types in the hierarchy (i.e., leaf nodes), compared with parent populations that aggregate multiple subtypes. Similar patterns were observed in the CITE-Seq dataset.

**Figure 1.**
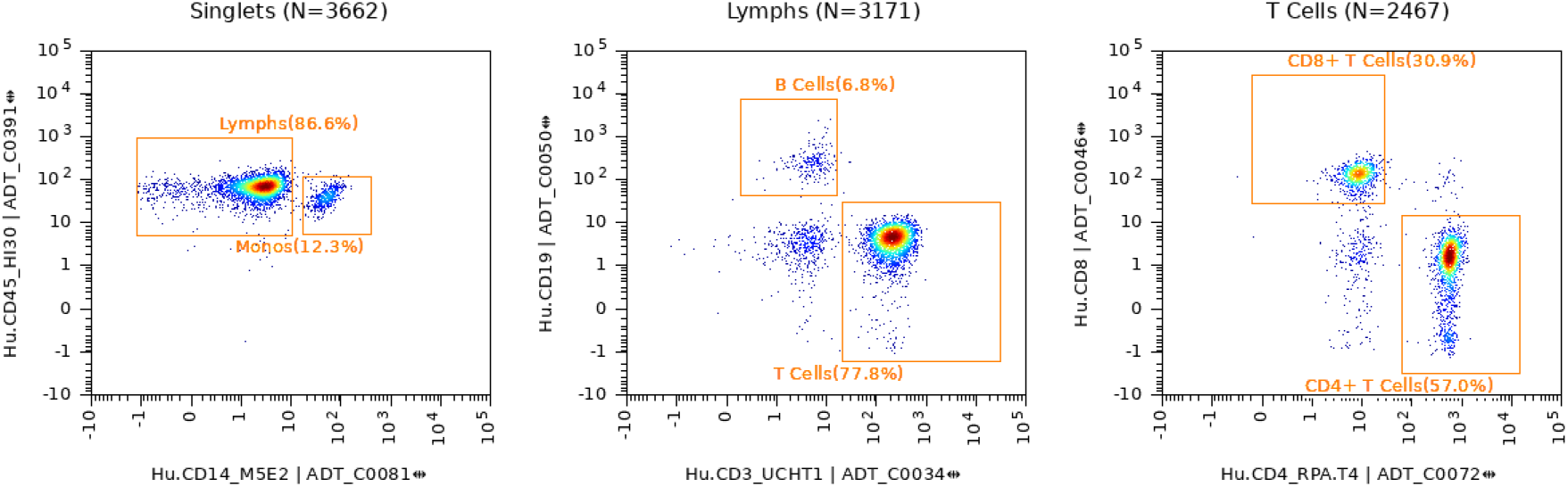
Typical gating plots showing CD14 and CD45 expression used to identify *Monocytes* and *Lymphocytes*, followed by CD3 and CD19 expression in *Lymphocytes* used to identify *T Cells* and *B Cells*, followed by CD4 and CD8 expression in *T Cells* used to identify *CD4+ T Cells* and *CD8+ T Cells*. In a plot, each point represents a cell, the color (red for high and blue for low) represents the density of cell in a particular region of the plot.

**Figure 2.**
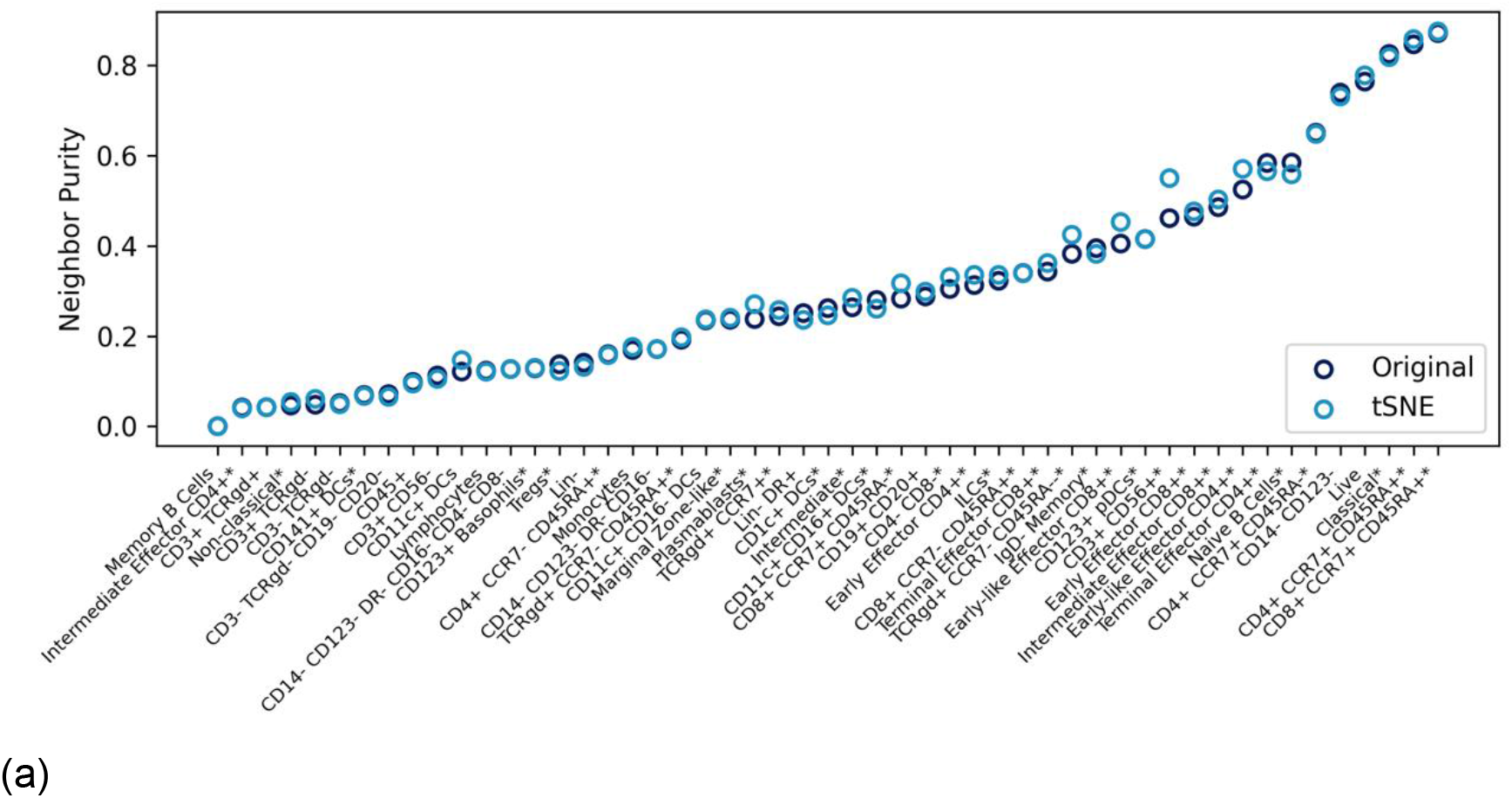
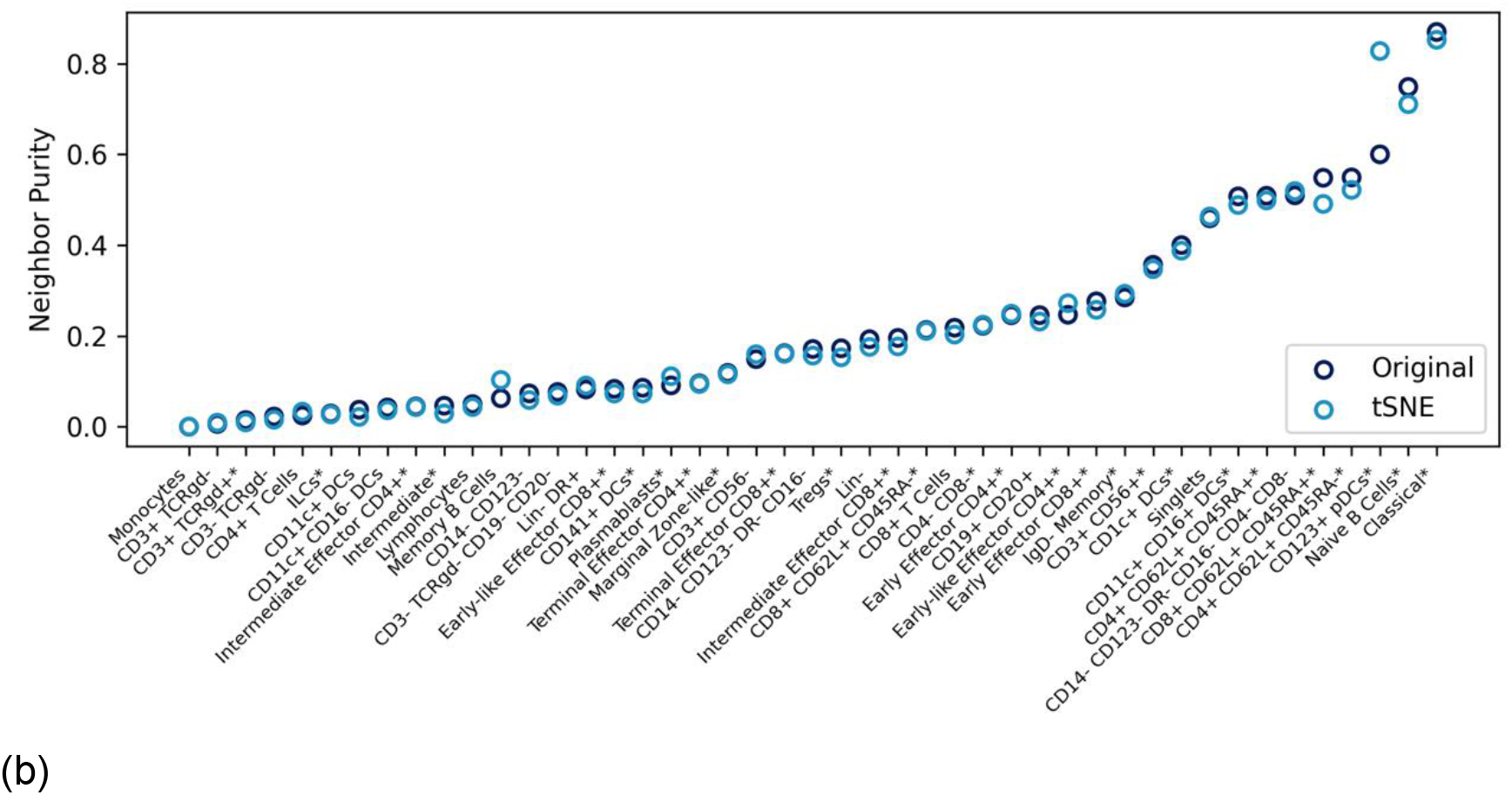
Neighborhood purity for each cell type a) Cytometry dataset b) CITE-Seq dataset. Color indicates if the neighbors were assessed in the original high dimensional space or in the projection space. Marker shape indicates if a cell type is a leaf node (i.e. child cell type) or a parent cell type.

#### 2.2.3 t-SNE correctly reflects the high dimensional space in the projection space

Examining neighborhood purity in the projection space (Figure 3) showed that purity values for each cell type closely match those observed in the original high-dimensional space. This indicates that t-SNE successfully achieves its primary objective—preserving local neighborhood structure during dimensionality reduction. Accordingly, the overlap between cell types observed in the projection space (Figures 4a and 4c) reflects the intrinsic structure of the data rather than a methodological limitation of t-SNE.

**Figure 3.**
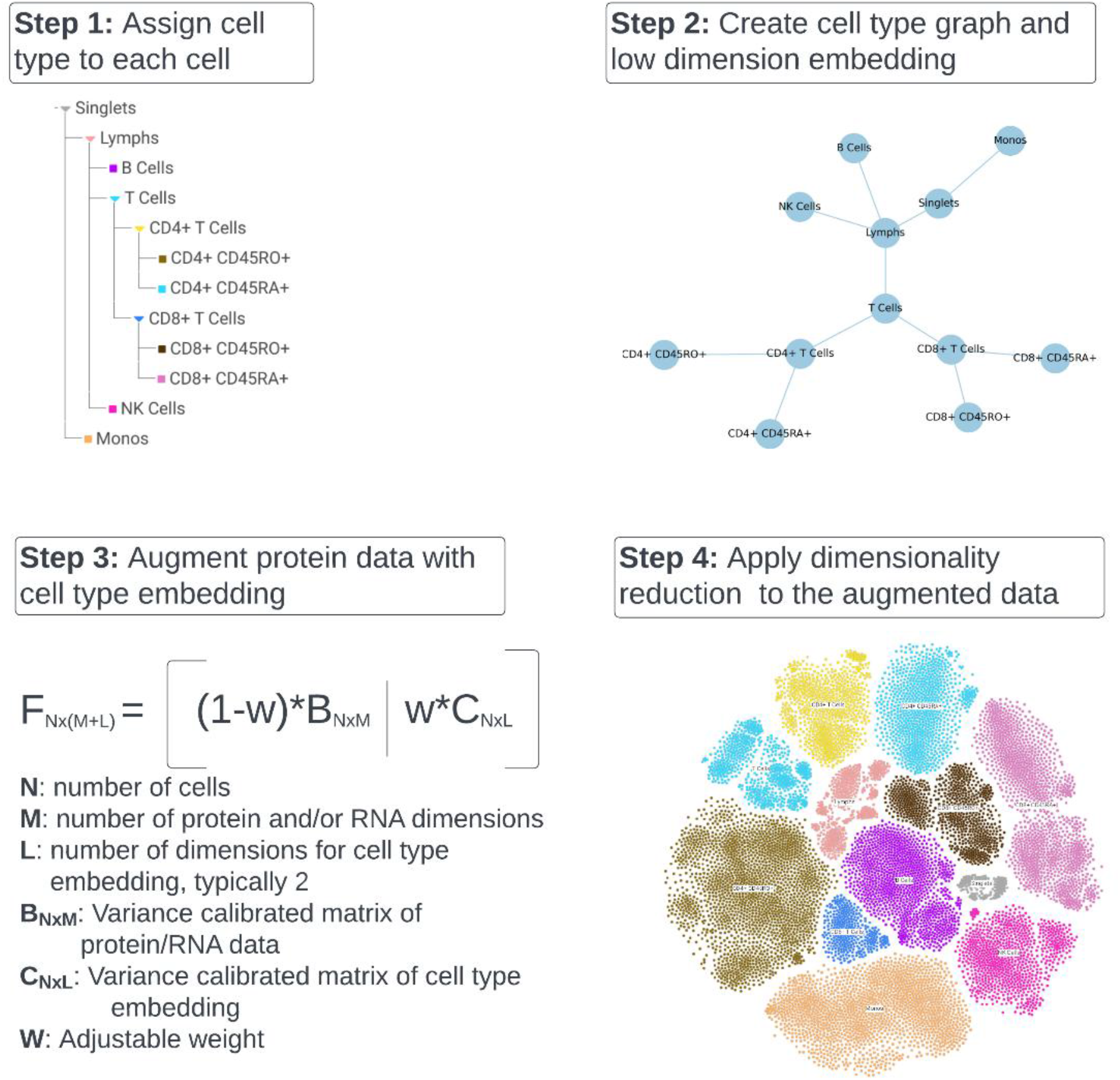
Steps to creating Cell Type Weighted Dimensionality Reduction: **Step 1:** Cell type is assigned to each cell typically using gating in a hierarchical order, with a child cell type taking precedence over the parent; **Step 2:** Create a cell type graph, compute distances (shortest path) between cell types from the graph and embed the graph in low (typically two) dimensional space, **Step 3:** Augment protein and RNA data for each cell with coordinates for each cell type from embedding from Step 2, **Step 4:** Apply dimensionality reduction methods like t-SNE or UMAP to the augmented dataset.

**Figure 4.**
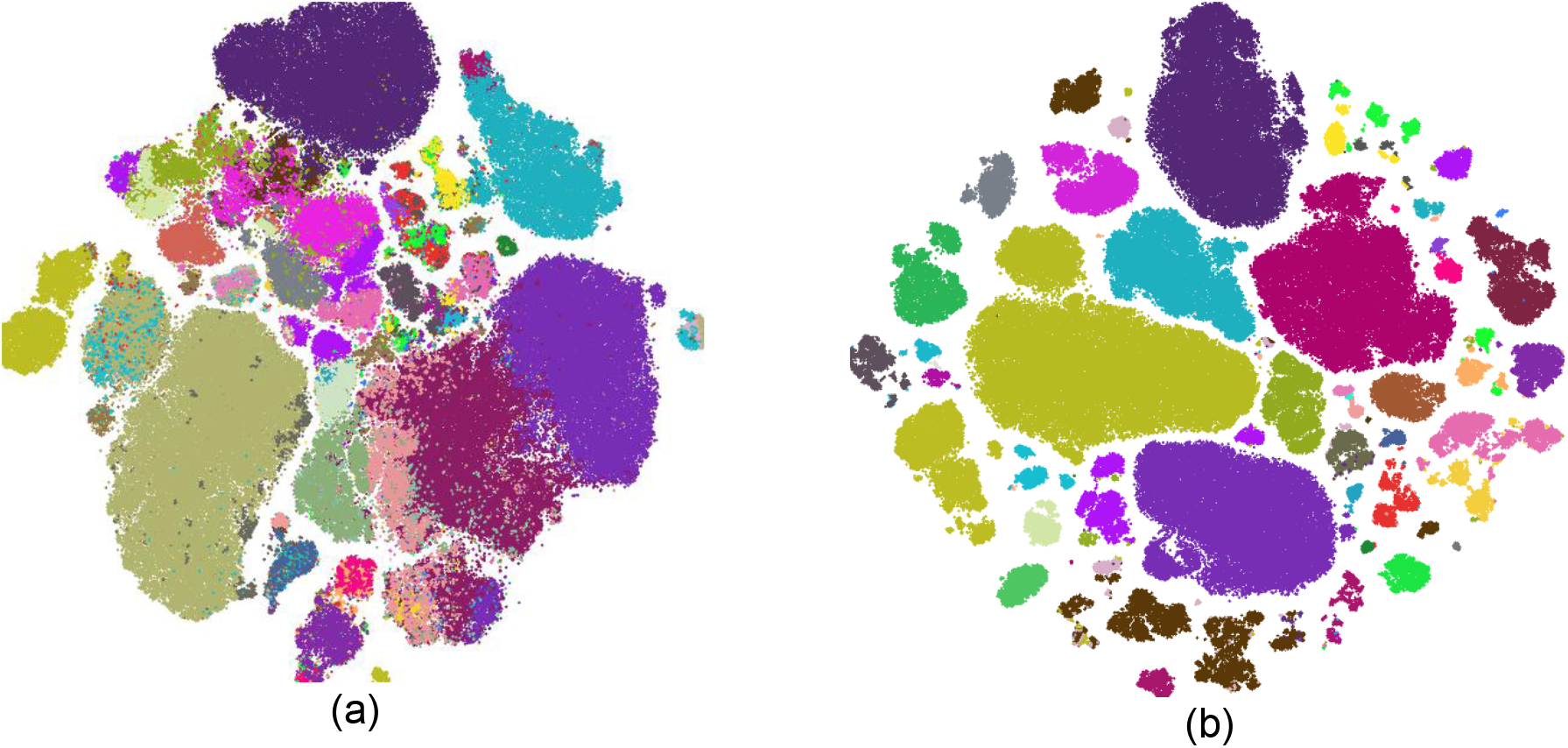
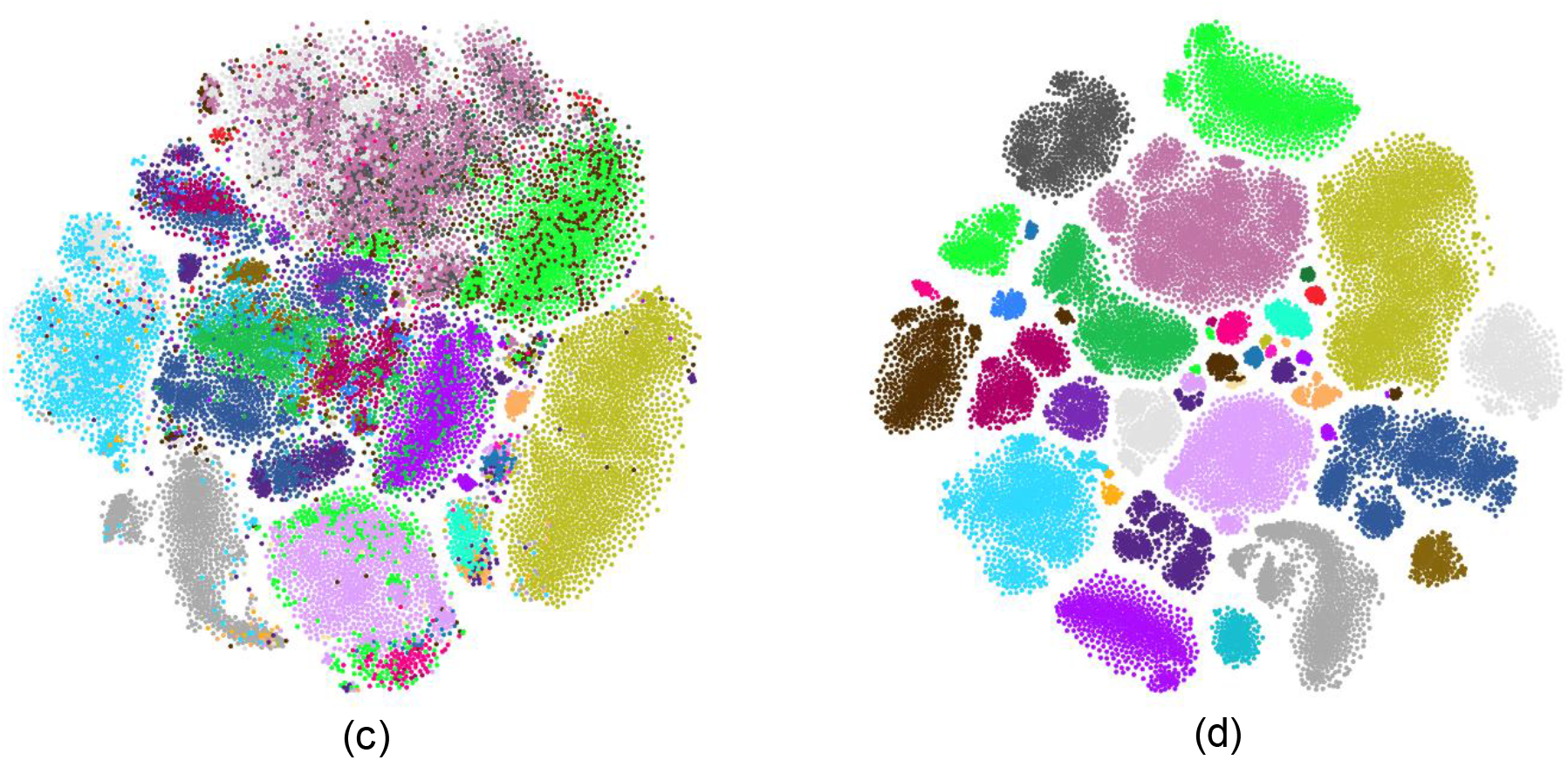
Results from t-SNE and Cell Type Weighted Dimensionality Reduction t-SNE applied to single cell data for PBMCs collected by flow cytometry (27 dimensions with 373115 cells, first row) and CITE-Seq (10678 cells and 153 dimensions of protein expression, second row); a) and c) t-SNE with no weighting for cell types, b) and d) Cell Type Weighted Dimensionality Reduction t-SNE with weight set to 0.5

These observations indicate that reducing cell-type overlap in the projection space requires incorporating cell-type information directly into the computation of each cell’s neighborhood.

### 2.3 Cell-type weighted dimensionality reduction delineates cell types in the projection space

Most dimensionality-reduction methods determine whether two points (cells) are neighbors based on their separation in the original high-dimensional space. The resulting projections then attempt to preserve these neighborhood relationships as faithfully as possible. For example, t-SNE computes Euclidean distances in the high-dimensional space, converts them into probabilistic neighborhood affinities, and uses an optimization procedure to match these probabilities in the low-dimensional embedding. Although methodologically distinct, UMAP follows a similar principle: it identifies nearby cells using a distance metric—typically Manhattan distance—to construct a weighted graph that reflects local connectivity and then optimizes a low-dimensional representation that preserves this graph structure. Both methods allow adjustment of distance metrics and neighborhood-related parameters to optimize projections for different datasets.

The primary input to dimensionality-reduction (DR) methods is the high-dimensional feature matrix, which does not explicitly encode cell-type information. Although cell types are derived from the same underlying data, the procedures for assigning cell identities differ fundamentally from those used to compute distances—and thus neighborhoods—between cells. Cell typing relies on a hierarchical lineage structure, typically using one or two markers at each decision point, whereas distance calculations treat all dimensions with equal weighting and provide no mechanism to incorporate hierarchical relationships. We therefore reasoned that integrating cell-type information directly into the distance computation would naturally reduce the likelihood of cell types overlapping in the resulting projection space.

#### 2.3.1 Cell-type weighted dimensionality reduction method

CWDR incorporates cell type based weighting mechanism into the distance computation prior to applying DR methods like t-SNE and UMAP. The process followed by CWDR is outlined below:

1. **Cell type assignment:** Cell type is assigned to each cell ordered by the hierarchy such that children take precedence over parents. For example, in the hierarchy shown in Figure 3, only those cells that have not been assigned to *CD4+ CD45RA+* or *CD4+ CD45RO+* are assigned to *CD4+ T Cells*.
2. **Graph construction for the cell-type hierarchy:** An acyclic graph is constructed for the cell type hierarchy (Figure 3). Each child cell type is connected directly to its parent (e.g., T cells connected to Lymphocytes). By default, each edge is assigned a weight of 1.0.
  a. All pairwise distances (length of shortest path) between the cell types are computed from the graph adjacency matrix. In the simplest form, the distance between two cell types corresponds to the minimum number of edges connecting them. For example, the distance between T cells and Lymphocytes is 1, while the distance between CD4^+^ T cells and Lymphocytes is 2. Several intuitive extensions to this metric are possible. One such modification accounts for the relative abundance of each cell type by weighting the distance between two connected cell types by the square root of the sum of their cell-type frequencies such that more space is available in the projection for cell types with more cells. This adjustment is made directly to the adjacency matrix, ensuring that all computed pairwise distances automatically reflect frequency-based weighting.
  b. A low-dimensional (typically two-dimensional) embedding is generated from the pairwise distance matrix produced in Step 2a. This embedding may be produced using t-SNE^1^, multidimensional scaling (MDS)^6^, or graph-layout approaches such as force-directed methods^7-9^. Importantly, this embedding depends solely on the cell-type hierarchy and is independent of protein or RNA expression data. By default, t-SNE is used for this step.
3. **Augmentation of single-cell data with cell-type coordinates:** For each cell, the original high-dimensional biological feature set (protein and/or RNA expression) is variance-normalized and augmented with the corresponding cell-type coordinates from Step 4. These coordinates constitute the cell-type dimensions (CTDs). Because CTDs arise from graph-based embedding, their coordinate system is arbitrary and unrelated to the biological dimensions (BDs). Variance normalization (division by the sum of feature standard deviations) places BDs and CTDs on comparable scales. An adjustable weight *w* then governs the relative contribution of CTDs to BDs when the combined feature matrix is constructed.
4. **Dimensionality-reduction of the augmented feature matrix:** In the last step, dimensionality reduction is applied to the augmented matrix (F_Nx(M+2)_ in Figure 3), producing a projection that incorporates both molecular measurements and lineage-informed structure.

An alternative, but equivalent, strategy for incorporating cell-type information into dimensionality reduction is to intervene directly during the similarity computation (as in t-SNE) or nearest-neighbor estimation (as in UMAP) in Step 4. Because these steps rely on pairwise distances between cells, one could define the distance between two cells as a weighted average of their biological-dimension (BD) distance and the cell-type distance derived from Step 3. From an implementation standpoint, however, generating cell-type dimensions (CTDs) first and augmenting the normalized BD matrix was a more practical approach. This design also enables the augmented matrix—combining normalized BDs and CTDs—to be used with any dimensionality-reduction method without requiring method-specific modifications.

#### 2.3.2 Cell Type Weighted Dimensionality Reduction provides better visual separation between cell types

Figure 4 illustrates t-SNE and Cell Type Weighted Dimensionality Reduction t-SNE applied to a PBMC (Peripheral Blood Mononuclear Cells) measure by both cytometry and CITE-Seq. As is clear from Figure 4b, cells belonging to distinct types are clearly visually separated. Additional sub-clusters within each type (e.g., *CD4+ Cells*) are visible. As discussed below, these sub-clusters are driven by protein dimensions that are unrelated to cell typing.

#### 2.3.3 Weighting provides a convenient mechanism to adjust the relative importance of cell type and marker dimensions

Figure 5 illustrates the results of applying Cell Type Weighted Dimensionality Reduction t-SNE across weighting levels ranging from 0.0 to 0.75. As expected, increasing the cell-type weight produces progressively tighter clusters of cells belonging to each respective cell type.

**Figure 5.**
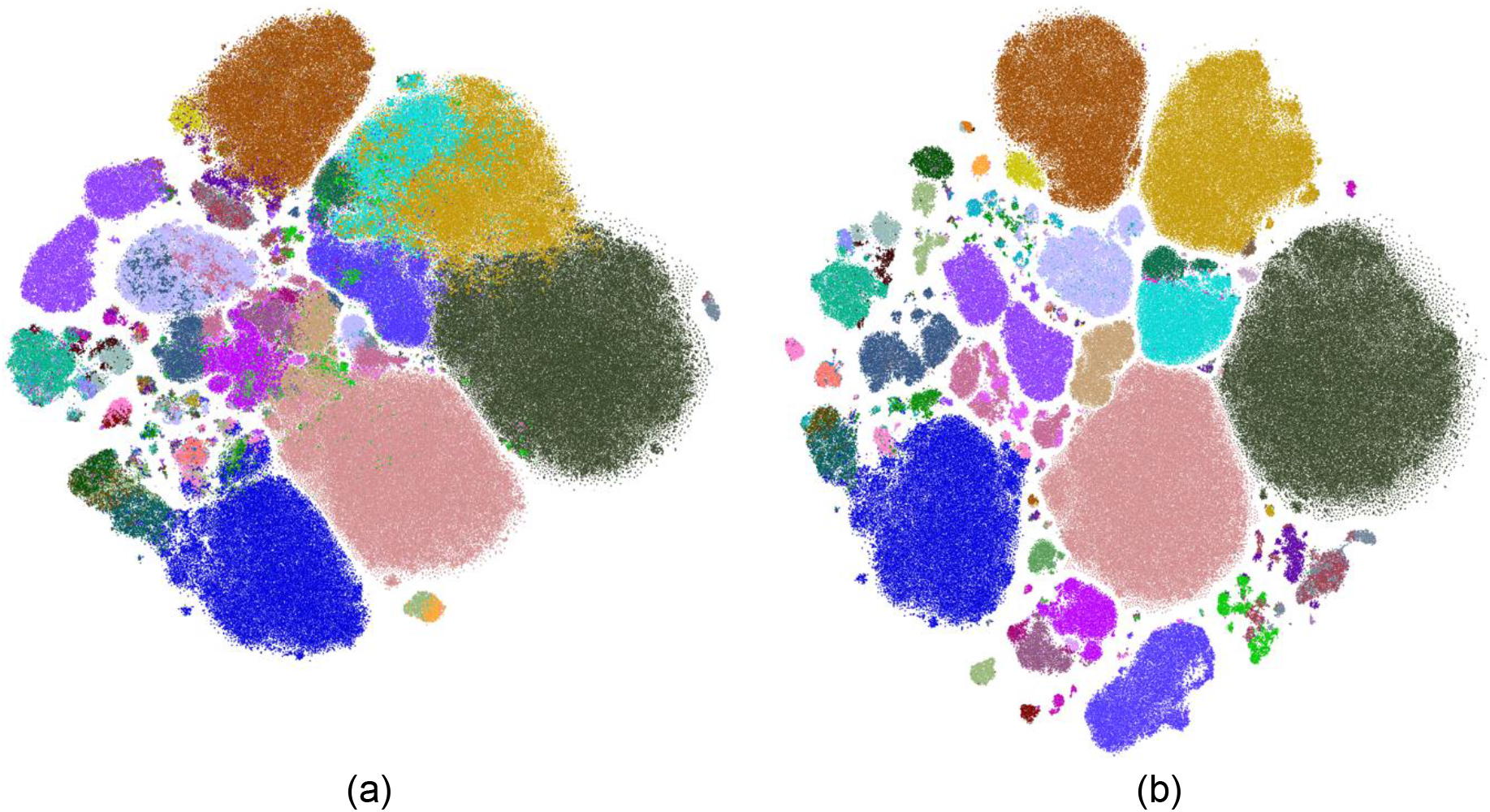
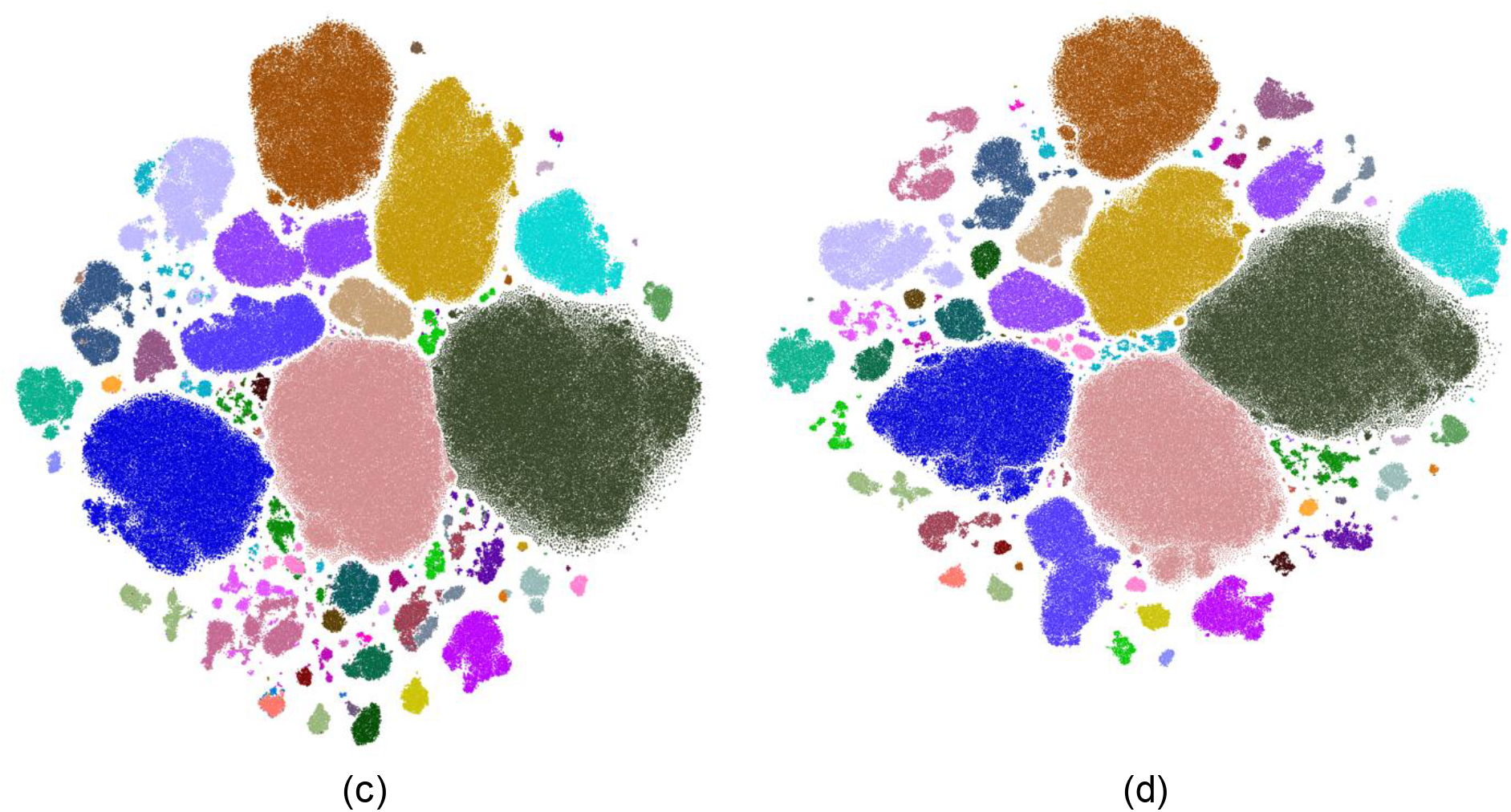
Cell Type Weighted Dimensionality Reduction t-SNE at increasing levels of cell type weighting applied to flow cytometry data: (a) 0.0, (b) 0.25, (c) 0.5, and (d) 0.75.

The overlap between cell types is reduced by CWDR, as confirmed by the neighborhoodpurity metric for each population. Figure 6 compares purity values computed using the weighted data (i.e., the FN × (M + 2) matrix in Figure 3) with those derived from the CWDR projections. In all cases, purity values are higher for the weighted method than for the original data. Notably, in several instances the purity observed in the projection space exceeds that of the weighted data itself. We hypothesize that this occurs because tSNE more easily preserves local neighborhoods while allowing cells from different types to drift farther apart in the lowdimensional embedding.

**Figure 6.**
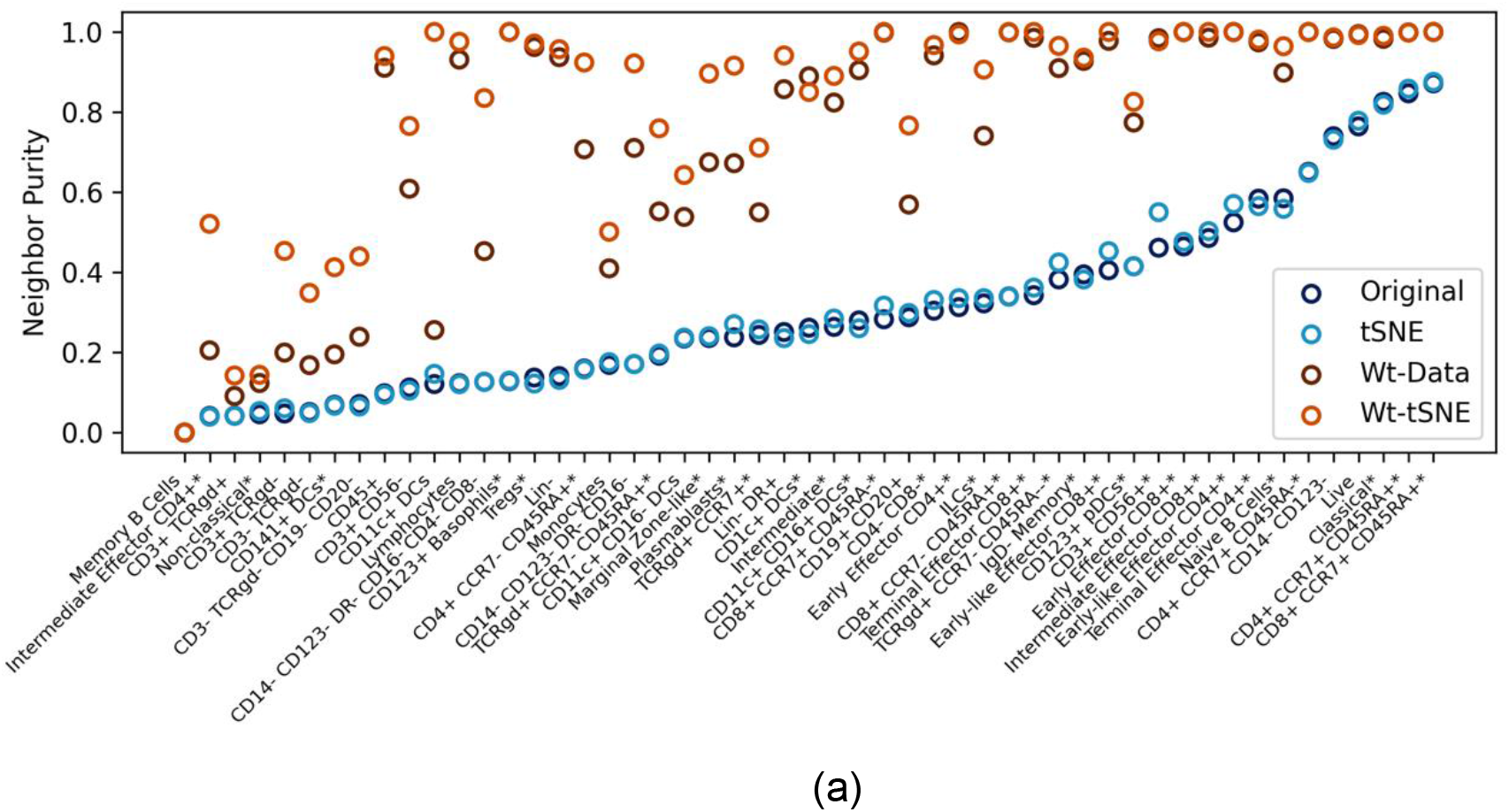
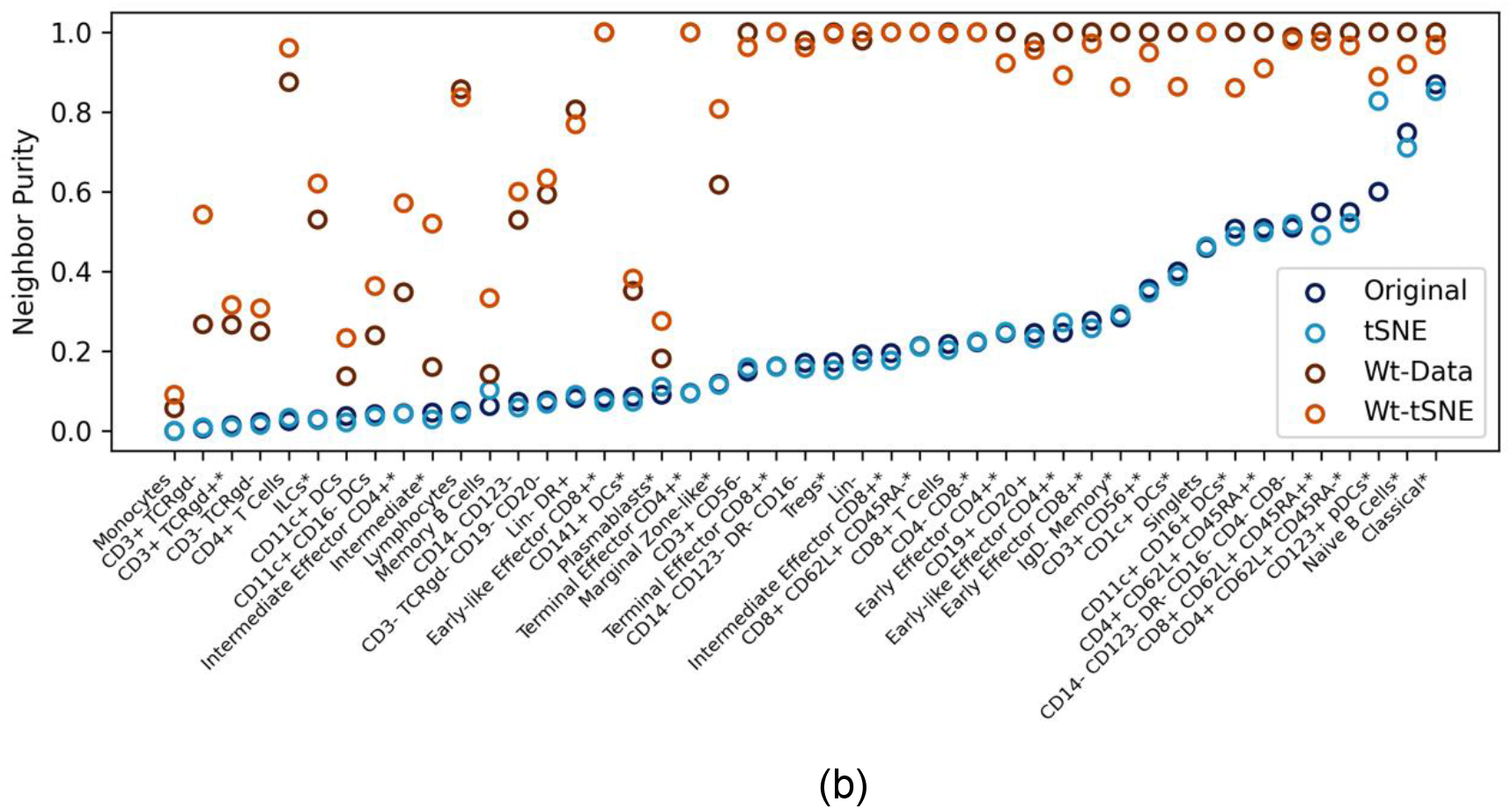
Neighborhood purity for each cell type the original data and weighted data a) Cytometry dataset b) CITE-Seq dataset. Color indicates if the neighbors were assessed in the original high dimensional space or in the project space. Marker shape indicates if a cell type is a leaf node (i.e., child cell types) or a parent cell type.

#### 2.3.4 Sub-clusters within a cell type are driven by other marker expression patterns

Cells belonging to the same cell type share identical CTD values; accordingly, CTDs do not contribute to any separation observed within a cell type. Any intra-type separation (i.e., sub-clustering) is therefore driven entirely by biological dimensions (BDs). In contrast, CTDs enhance dissimilarity between cells of different types while still incorporating BD information. As such, CTDs provide an adjustable counterbalance to BD-driven similarity across distinct cell types. Within a given cell type, dimensionality-reduction methods such as t-SNE continue to highlight sub-clusters arising from heterogeneous expression of markers not used for cell-type assignment.

For example, in Figure 7, CD3^+^CD56^+^ (NKT) cells exhibit multiple sub-clusters. Overlaying marker expression on these projections shows that CD4, CD45RA, and TCR-Vγ9 are among the primary drivers of this intra-type heterogeneity.

**Figure 7.**
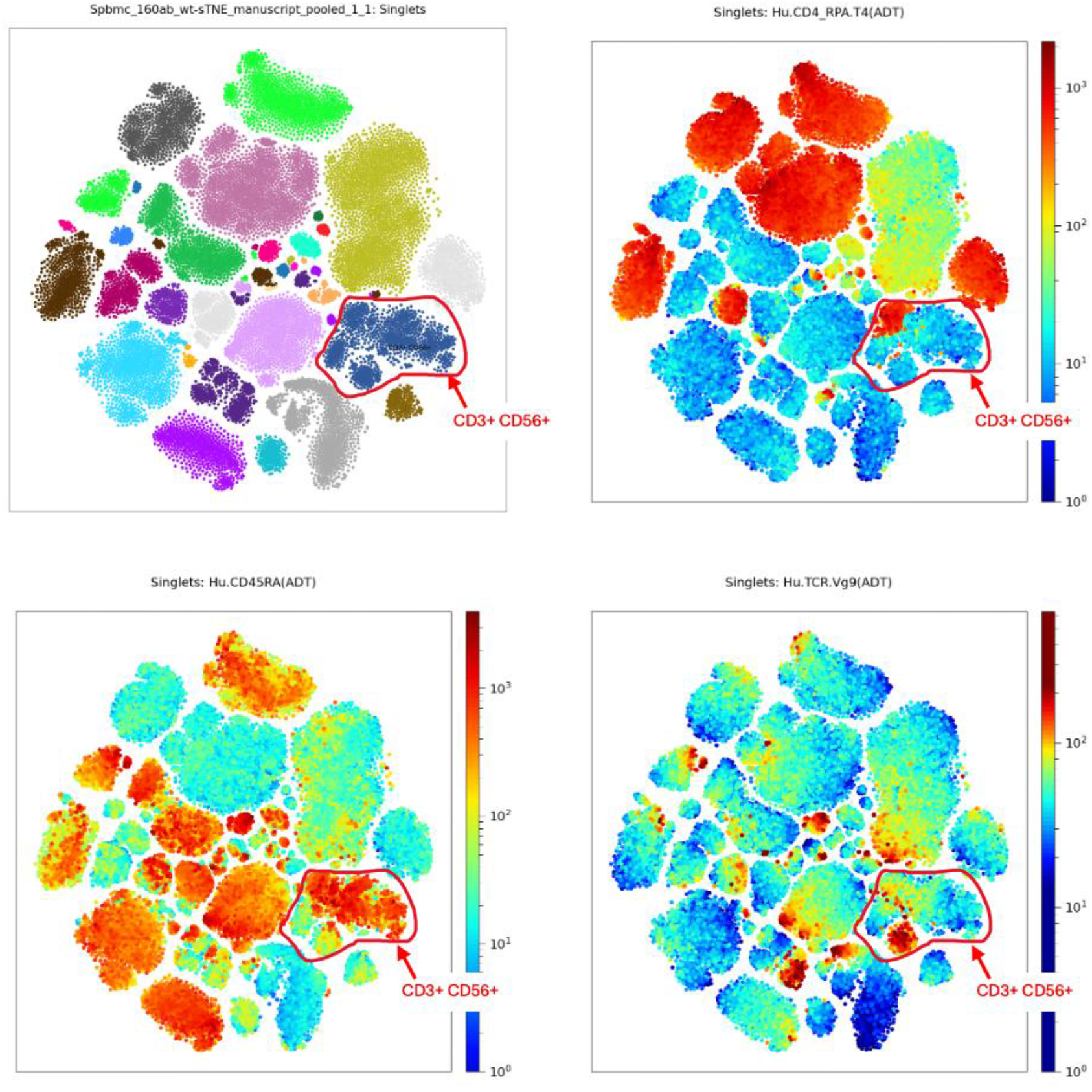
Within each cell type markers that were not used for cell type determination can drive the sub-clusters. For example, *CD3+ CD56+* cells display sub-clusters in (a). Examination of other markers like CD4 (b), CD45RA(c) and TCR.Vg9(d) show that these sub-clusters within *CD3+ CD56+* cells have distinct expression patterns.

**Figure 8.**
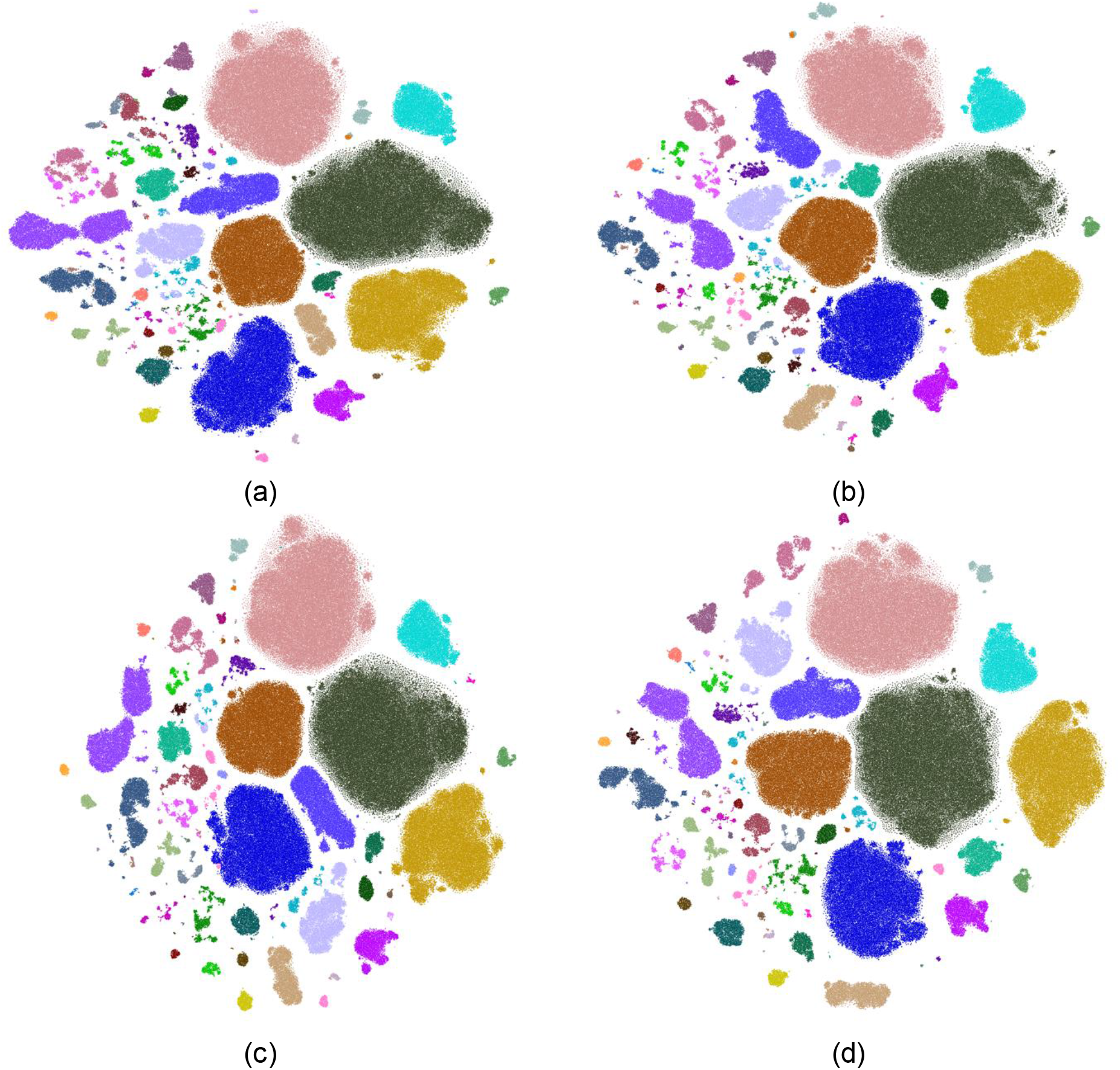
Cell Type Weighted t-SNE **r**esults when using different methods for computing CTDs a) t-SNE, b) MDS, c) Spring Layout d) Kamada-Kawai layout.

#### 2.3.5 Choice of embedding method for cell type graph has minimal impact on weighted t-SNE results

As noted in Section 2.3.1, multiple methods can be used to compute CTDs from the cell-type graph. The default approach applies t-SNE to the pairwise distances between cell types, where distances are defined as shortest-path lengths in the graph. Alternatively, methods such as multidimensional scaling (MDS) or graph-layout algorithms, including Spring or Kamada–Kawai layouts, may be used. We evaluated these approaches and found that all performed comparably. Figure 6 presents Cell Type Weighted t-SNE results generated using four different CTD-embedding methods.

## 3. Limitations of Study

The primary objective of this work is to develop a dimensionality-reduction method that provides an effective visual framework for displaying protein and RNA expression within the context of established cell types. As described above, we achieve this by modifying the input data used to compute inter-cell distances, and thus the neighborhood relationships that drive the projection. Consequently, the resulting embeddings do not reflect the conventional notion of proximity in marker space. Unlike standard methods such as t-SNE, CWDR is not intended to represent neighborhoods in the original feature space. Therefore, the projections it generates should not be interpreted in that manner.

## 4. Discussion

A primary function of dimensionality reduction is to facilitate the visualization of high-dimensional data. In single-cell biology, such visualization is most informative when interpreted within established biological frameworks, particularly cell types. We showed that standard methods such as t-SNE cannot fully separate diverse cell types in projection space— not due to methodological shortcomings, but because cells from different types can reside in close proximity in the original high-dimensional space when distances are computed using unbiased metrics (e.g., Euclidean distance with equal weights across features). Consequently, dimensionality-reduction methods cannot be expected to disentangle cell types in low-dimensional space unless cell-type information is explicitly incorporated.

The method we introduce provides such a mechanism by integrating cell-type information directly into the dimensionality-reduction process. This is achieved by augmenting the single-cell feature matrix with graph-based embeddings of the cell-type hierarchy. The framework includes a tunable parameter that modulates the relative influence of biological features and cell-type structure. As demonstrated, this approach accomplishes two key goals: (1) improving the spatial separation of distinct cell types, and (2) preserving and revealing intra-type sub-clusters driven by heterogeneous marker expression.

A potential criticism of our approach is that incorporating cell typing may bias the projection, potentially giving the misleading impression that cell-type separation in the embedding reflects actual distances in the original feature space. Prior work^9,10^ has emphasized that while dimensionality-reduction methods such as t-SNE and UMAP reasonably preserve local neighborhoods, they do not reliably maintain global structure. Accordingly, these approaches— including ours—should be used primarily as tools for visual exploration rather than quantitative interpretation of global geometry. Our method adheres to this principle and extends it by enabling visualization that aligns more closely with known cell-lineage relationships.

Beyond visualization, our findings highlight a broader conceptual issue: conventional distance metrics computed from high-dimensional protein data do not necessarily align with biologically defined cell types. Because cell-type assignments depend on hierarchical marker usage rather than uniform weighting across all features, cells from distinct lineages may appear close in the original feature space despite being biologically distinct. This mismatch becomes increasingly pronounced as the depth of the lineage hierarchy grows. We anticipate that this disconnect also affects clustering methods^12,13^, many of which rely fundamentally on neighborhood definitions derived from these same distance metrics. While further exploration is beyond the scope of this manuscript, future work may include developing automated approaches for cell typing that address this discrepancy more directly.

## 5. Acknowledgements

All the work detailed in this manuscript was funded by Revvity, Inc. The authors would also like to thank David Soper and Rebecca Nickle for their valuable feedback on the manuscript.

## References

1. Visualizing Data using t-SNE, Laurens van der Maaten, Geoffrey Hinton, Journal of Machine Learning Research 9 (2008) 2579–2605

2. UMAP: Uniform Manifold Approximation and Projection, Leland McInnes, John Healy, Nathaniel Saul, Lukas, Großberger Journal of Open-Source Software, 3(29), 861, 10.21105/joss.00861

3. Multiomics Analysis Software (MAS): Cloud software for the analysis of single-cell proteogenomics data. https://www.biolegend.com/en-us/totalseq/mas

4. CytoScribe−: Cloud software for flow-cytometry data analysis. https://www.biolegend.com/en-us/cytoscribe

5. Faiss: A library for efficient similarity search. https://engineering.fb.com/2017/03/29/data-infrastructure/faiss-a-library-for-efficient-similarity-search/

6. Multidimensional scaling by optimizing goodness of fit to a nonmetric hypothesis, Kruskal, J. B., Psychometrika. 29 (1): 1–27, 1964

7. Graph Drawing by Force-Directed Placement, Fruchterman, Thomas M. J.; Reingold, Edward M. (1991), Software: Practice and Experience, Wiley, 21 (11): 1129–1164, doi:10.1002/spe.4380211102

8. An algorithm for drawing general undirected graphs, Kamada, Tomihisa; Kawai, Satoru (1989), Information Processing Letters, Elsevier, 31 (1): 7–15, doi:10.1016/0020-0190(89)90102-6

9. Spring Embedders and Force-Directed Graph Drawing Algorithms, Kobourov, Stephen G. (2012), https://arxiv.org/abs/1201.3011

10. Towards a comprehensive evaluation of dimension reduction methods for transcriptomic data visualization, Huang, H., Wang, Y., Rudin, C. et al., Commun Biol 5, 719 (2022). 10.1038/s42003-022-03628-x

11. Understanding How Dimension Reduction Tools Work: An Empirical Approach to Deciphering t-SNE, UMAP, TriMap, and PaCMAP for Data Visualization, Yingfan Wang, Haiyang Huang, Cynthia Rudin, and Yaron Shaposhnik, Journal of Machine Learning Research 22 (2021) 1–73

12. Leiden: From Louvain to Leiden: guaranteeing well-connected communities, V. A. Traag, L. Waltman & N. J. van Eck; https://arxiv.org/abs/1810.08473

13. PARC: Ultrafast and accurate clustering of phenotypic data of millions of single cells, Shobana V Stassen, Dickson M D Siu, Kelvin C M Lee, Joshua W K Ho, Hayden K H So, Kevin K Tsia; Bioinformatics, Volume 36, Issue 9, May 2020, Pages 2778–2786

