## Supplemental for "Cell Type Weighted Dimensionality Reduction"

### Supplementary Materials: Cell Type Weighted Dimensionality Reduction

A. Gating information for flow cytometry data

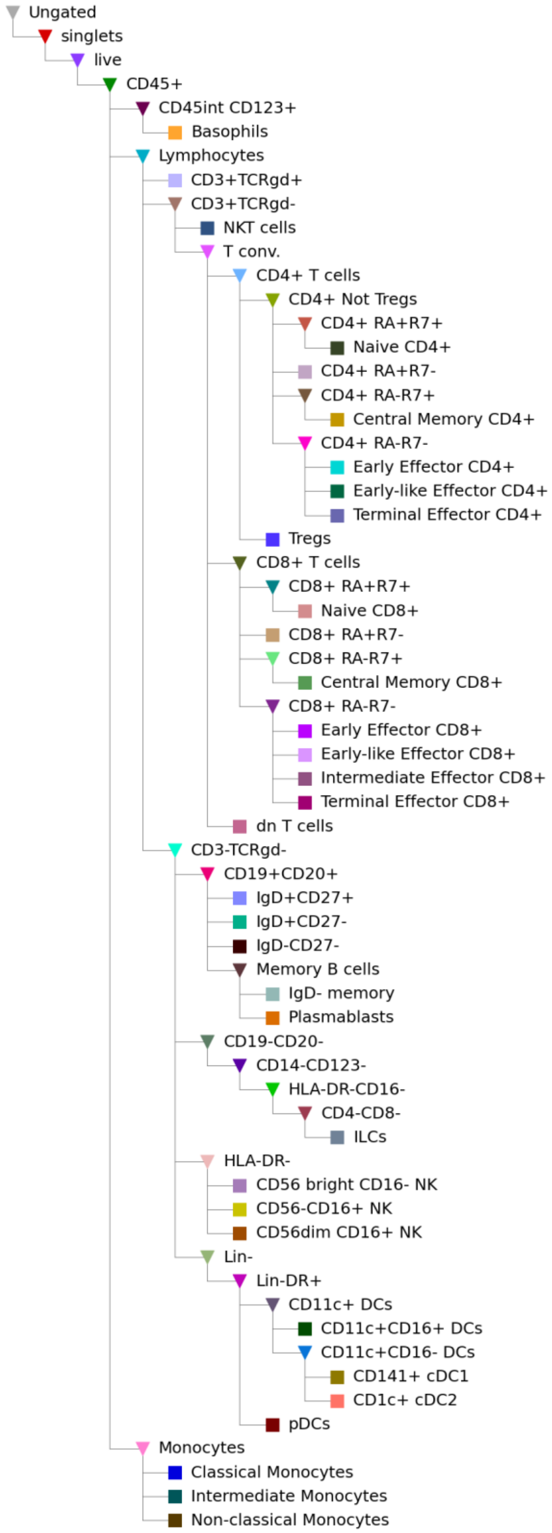

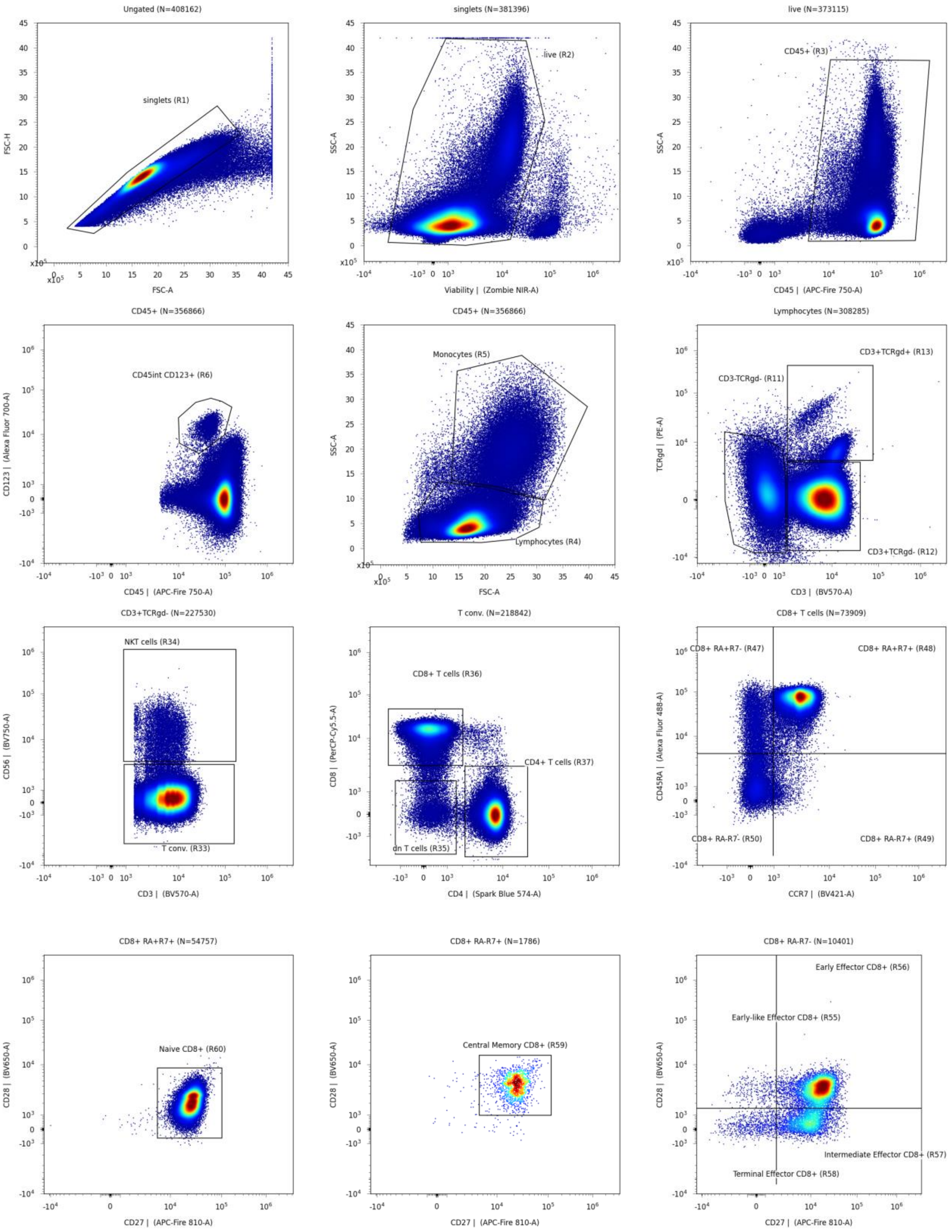
Ungated (N=408162)
singlets (R1)
FSC-H
FSC-A
singlets (N=381396)
live (R2)
SSC-A
Viability | (Zombie NIR-A)
live (N=373115)
CD45+ (R3)
SSC-A
CD45 | (APC-Fire 750-A)
CD45+ (N=356866)
CD45int CD123+ (R6)
CD123 | (Alexa Fluor 700-A)
CD45 | (APC-Fire 750-A)
CD45+ (N=356866)
Monocytes (R5)
Lymphocytes (R4)
SSC-A
FSC-A
Lymphocytes (N=308285)
CD3+TCRgd+ (R13)
CD3-TCRgd- (R11)
CD3+TCRgd- (R12)
TCRgd | (PE-A)
CD3 | (BV570-A)
CD3+TCRgd- (N=227530)
NKT cells (R34)
T conv. (R33)
CD56 | (BV750-A)
CD3 | (BV570-A)
T conv. (N=218842)
CD8+ T cells (R36)
CD4+ T cells (R37)
dn T cells (R35)
CD8 | (PerCP-Cy5.5-A)
CD4 | (Spark Blue 574-A)
CD8+ T cells (N=73909)
CD8+ RA+R7- (R47)
CD8+ RA+R7+ (R48)
CD8+ RA-R7- (R50)
CD8+ RA-R7+ (R49)
CD45RA | (Alexa Fluor 488-A)
CCR7 | (BV421-A)
CD8+ RA+R7+ (N=54757)
Naive CD8+ (R60)
CD28 | (BV650-A)
CD27 | (APC-Fire 810-A)
CD8+ RA-R7+ (N=1786)
Central Memory CD8+ (R59)
CD28 | (BV650-A)
CD27 | (APC-Fire 810-A)
CD8+ RA-R7- (N=10401)
Early Effector CD8+ (R56)
Early-like Effector CD8+ (R55)
Intermediate Effector CD8+ (R57)
Terminal Effector CD8+ (R58)
CD28 | (BV650-A)
CD27 | (APC-Fire 810-A)

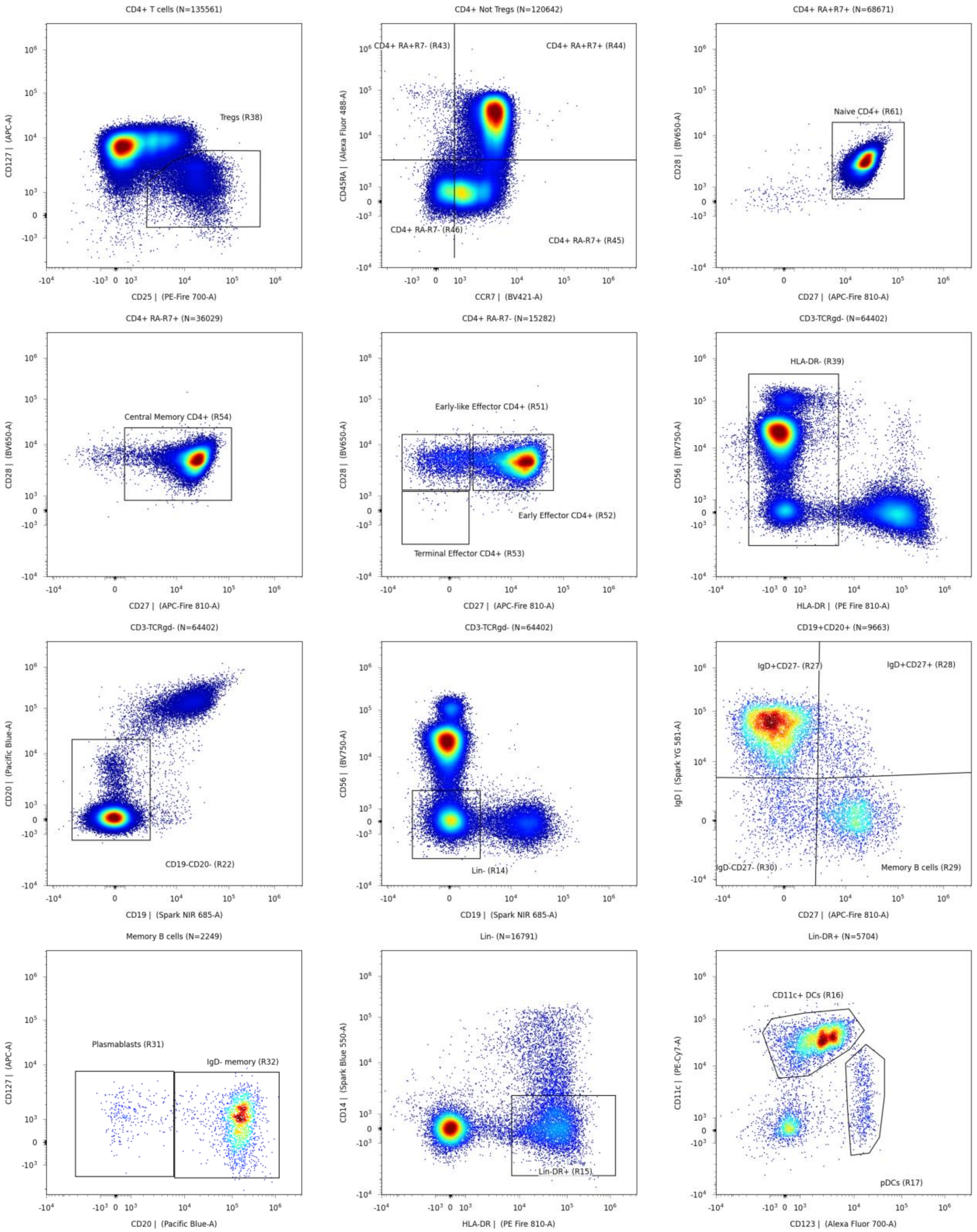
CD4+ T cells (N=135561)
Tregs (R38)
CD127 | (APC-A)
CD25 | (PE-Fire 700-A)
CD4+ Not Tregs (N=120642)
CD4+ RA+R7- (R43)
CD4+ RA+R7+ (R44)
CD4+ RA-R7- (R46)
CD4+ RA-R7+ (R45)
CD45RA | (Alexa Fluor 488-A)
CCR7 | (BV421-A)
CD4+ RA+R7+ (N=68671)
Naive CD4+ (R61)
CD28 | (BV650-A)
CD27 | (APC-Fire 810-A)
CD4+ RA-R7+ (N=36029)
Central Memory CD4+ (R54)
CD28 | (BV650-A)
CD27 | (APC-Fire 810-A)
CD4+ RA-R7- (N=15282)
Early-like Effector CD4+ (R51)
Early Effector CD4+ (R52)
Terminal Effector CD4+ (R53)
CD28 | (BV650-A)
CD27 | (APC-Fire 810-A)
CD3-TCRgd- (N=64402)
HLA-DR- (R39)
CD56 | (BV750-A)
HLA-DR | (PE Fire 810-A)
CD3-TCRgd- (N=64402)
CD19-CD20- (R22)
CD20 | (Pacific Blue-A)
CD19 | (Spark NIR 685-A)
CD3-TCRgd- (N=64402)
Lin- (R14)
CD56 | (BV750-A)
CD19 | (Spark NIR 685-A)
CD19+CD20+ (N=9663)
IgD+CD27- (R27)
IgD+CD27+ (R28)
IgD-CD27- (R30)
Memory B cells (R29)
IgD | (Spark YG 581-A)
CD27 | (APC-Fire 810-A)
Memory B cells (N=2249)
Plasmablasts (R31)
IgD- memory (R32)
CD127 | (APC-A)
CD20 | (Pacific Blue-A)
Lin- (N=16791)
Lin-DR+ (R15)
CD14 | (Spark Blue 550-A)
HLA-DR | (PE Fire 810-A)
Lin-DR+ (N=5704)
CD11c+ DCs (R16)
pDCs (R17)
CD11c | (PE-Cy7-A)
CD123 | (Alexa Fluor 700-A)

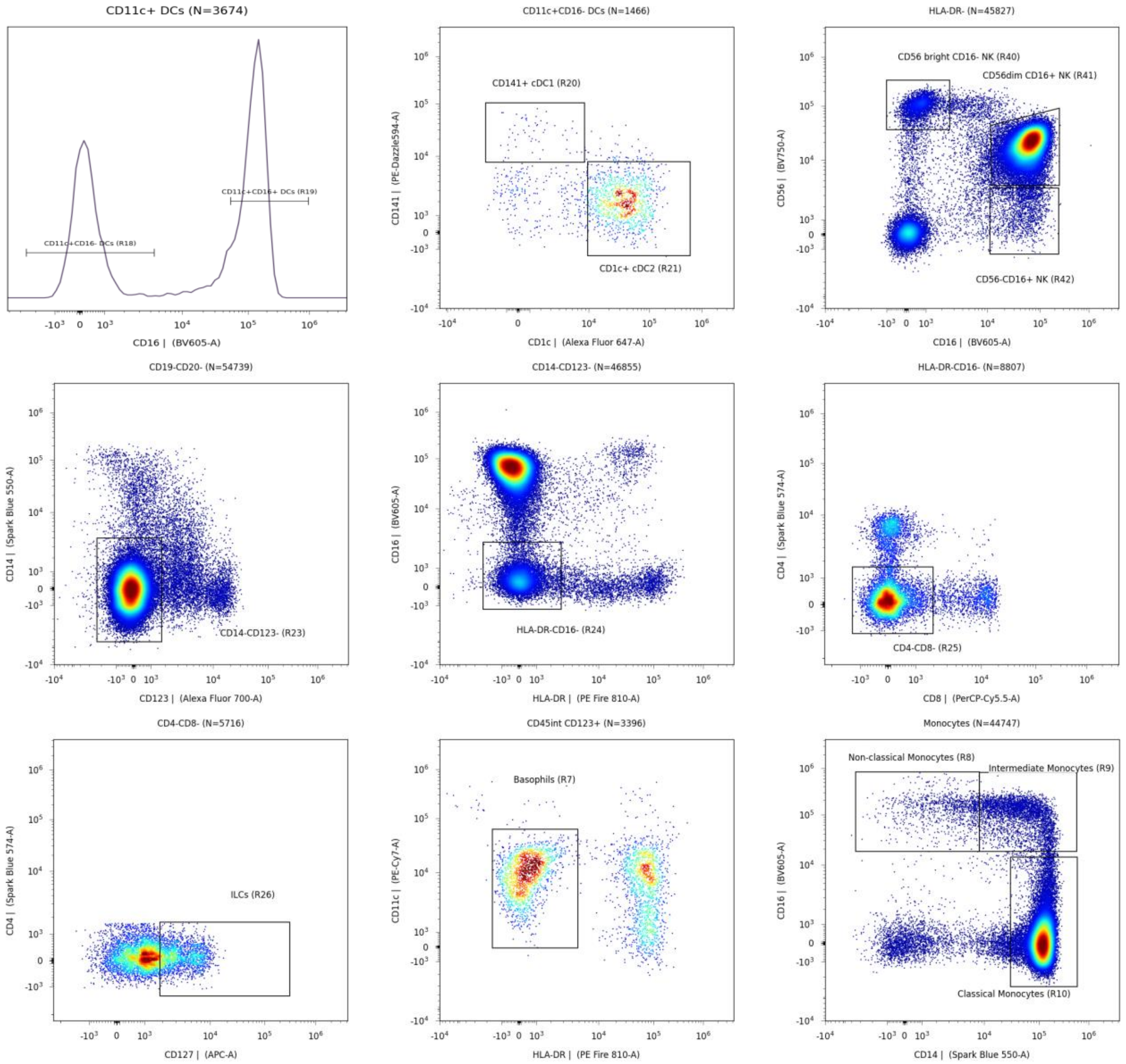


B. Gating information for CITE-Seq data

- Singlets (N=12456)
  - Monocytes (N=1623)
    - Non-classical (N=0)
    - Intermediate (N=13)
    - Classical (N=1608)
  - Lymphocytes (N=10079)
    - CD3- TCRgd- (N=2758)
      - Lin- (N=540)
        - Lin- DR+ (N=252)
          - CD123+ pDCs (N=24)
          - CD11c+ DCs (N=175)
            - CD11c+ CD16+ DCs (N=105)
            - CD11c+ CD16- DCs (N=63)
              - CD1c+ DCs (N=38)
              - CD141+ DCs (N=13)
      - CD3- TCRgd- CD19- CD20- (N=1575)
        - CD14- CD123- (N=1226)
          - CD14- CD123- DR- CD16- (N=1159)
            - CD14- CD123- DR- CD16- CD4- CD8- (N=881)
              - ILCs (N=36)
      - CD19+ CD20+ (N=1174)
        - Naive B Cells (N=824)
        - Marginal Zone-like (N=21)
        - Memory B Cells (N=92)
          - IgD- Memory (N=76)
          - Plasmablasts (N=8)
    - CD3+ TCRgd- (N=7237)
      - CD3+ CD56- (N=6321)
        - CD4- CD8- (N=360)
        - CD8+ T Cells (N=1977)
          - CD8+ CD62L+ CD45RA- (N=279)
          - CD8+ CD62L+ CD45RA+ (N=939)
          - CD8+ CD62L- CD45RA- (N=447)
            - Intermediate Effector CD8+ (N=104)
            - Terminal Effector CD8+ (N=87)
            - Early-like Effector CD8+ (N=46)
            - Early Effector CD8+ (N=210)
        - CD4+ T Cells (N=3776)
          - CD4+ CD62L+ CD45RA- (N=1927)
          - Tregs (N=277)
          - CD4+ CD62L+ CD45RA+ (N=1049)
          - CD4+ CD62L- CD45RA- (N=747)
            - Terminal Effector CD4+ (N=29)
            - Early-like Effector CD4+ (N=273)
            - Early Effector CD4+ (N=429)
            - Intermediate Effector CD4+ (N=16)
      - CD3+ CD56+ (N=897)
    - CD3+ TCRgd+ (N=12)

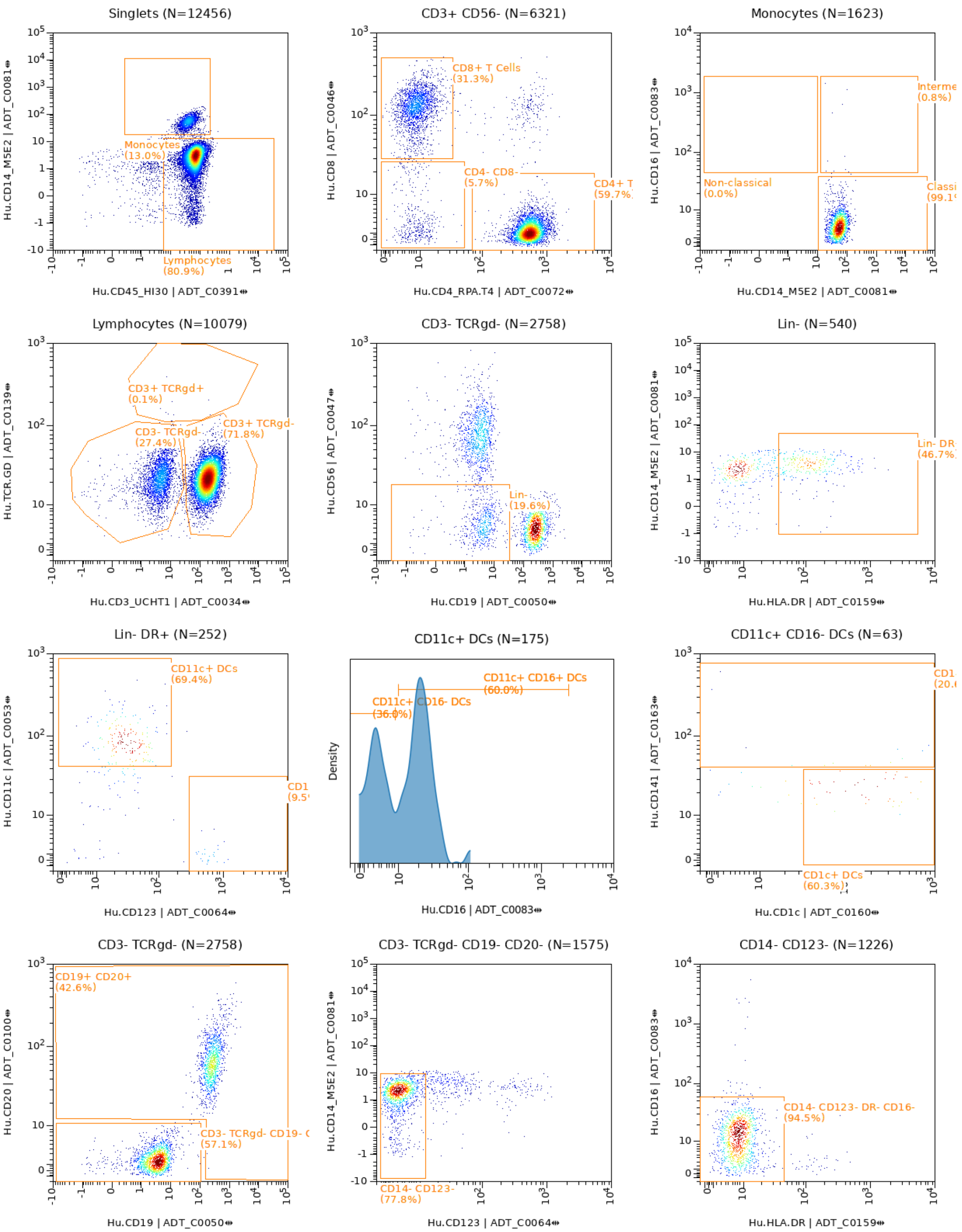
Singlets (N=12456)
Monocytes (13.0%)
Lymphocytes (80.9%)
Hu.CD14_M5E2 | ADT_C0081
Hu.CD45_HI30 | ADT_C0391
CD3+ CD56- (N=6321)
CD8+ T Cells (31.3%)
CD4- CD8- (5.7%)
CD4+ T (59.7%)
Hu.CD8 | ADT_C0046
Hu.CD4_RPA.T4 | ADT_C0072
Monocytes (N=1623)
Interme (0.8%)
Non-classical (0.0%)
Classi (99.1
Hu.CD16 | ADT_C0083
Hu.CD14_M5E2 | ADT_C0081
Lymphocytes (N=10079)
CD3+ TCRgd+ (0.1%)
CD3- TCRgd- (27.4%)
CD3+ TCRgd- (71.8%)
Hu.TCR.GD | ADT_C0139
Hu.CD3_UCHT1 | ADT_C0034
CD3- TCRgd- (N=2758)
Lin- (19.6%)
Hu.CD56 | ADT_C0047
Hu.CD19 | ADT_C0050
Lin- (N=540)
Lin- DR (46.7%)
Hu.CD14_M5E2 | ADT_C0081
Hu.HLA.DR | ADT_C0159
Lin- DR+ (N=252)
CD11c+ DCs (69.4%)
CD1 (9.5
Hu.CD11c | ADT_C0053
Hu.CD123 | ADT_C0064
CD11c+ DCs (N=175)
CD11c+ CD16+ DCs (60.0%)
CD11c+ CD16- DCs (36.0%)
Density
Hu.CD16 | ADT_C0083
CD11c+ CD16- DCs (N=63)
CD1 (20.
CD1c+ DCs (60.3%)
Hu.CD141 | ADT_C0163
Hu.CD1c | ADT_C0160
CD3- TCRgd- (N=2758)
CD19+ CD20+ (42.6%)
CD3- TCRgd- CD19- ( (57.1%)
Hu.CD20 | ADT_C0100
Hu.CD19 | ADT_C0050
CD3- TCRgd- CD19- CD20- (N=1575)
CD14- CD123- (77.8%)
Hu.CD14_M5E2 | ADT_C0081
Hu.CD123 | ADT_C0064
CD14- CD123- (N=1226)
CD14- CD123- DR- CD16- (94.5%)
Hu.CD16 | ADT_C0083
Hu.HLA.DR | ADT_C0159

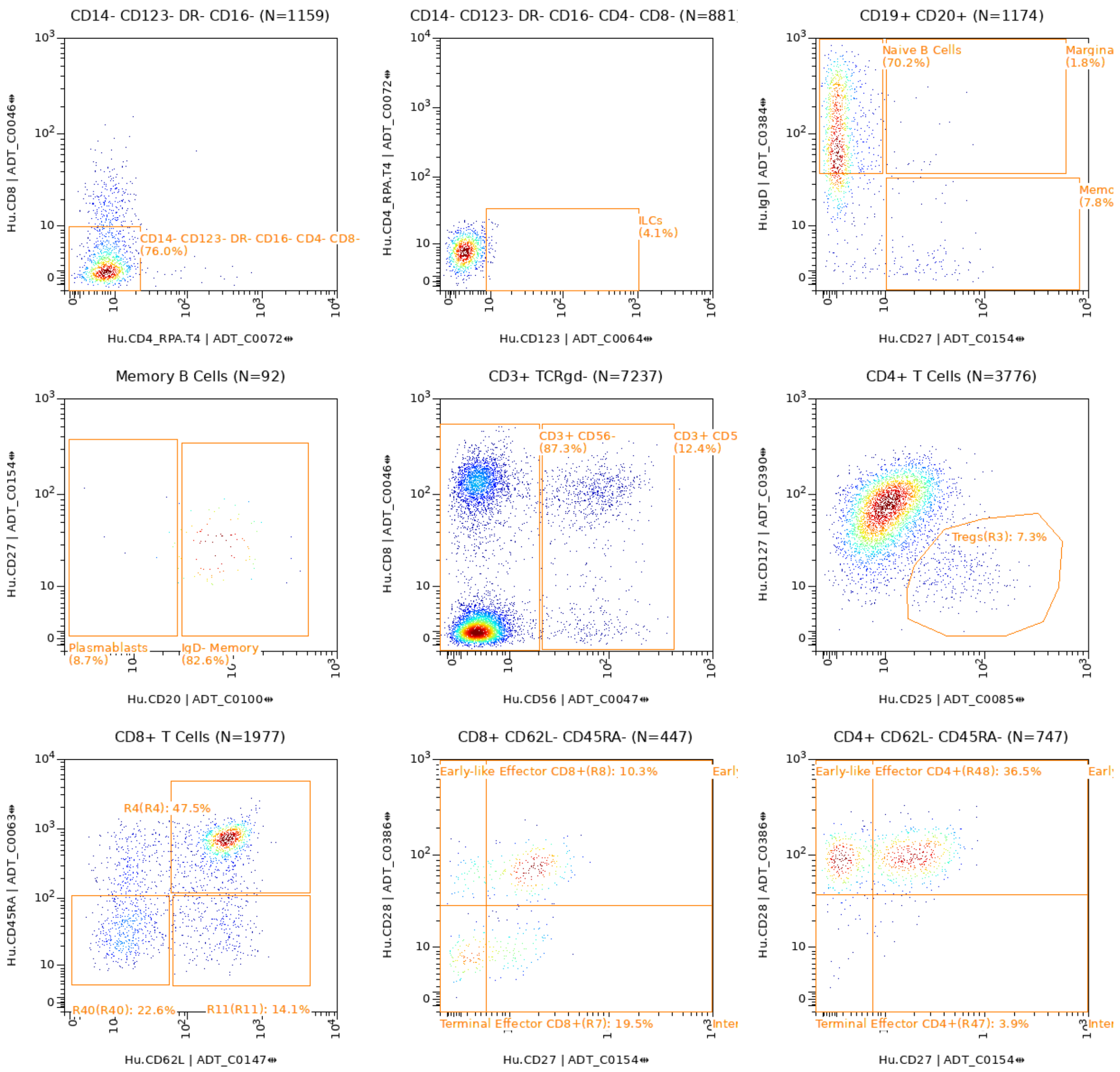
CD14- CD123- DR- CD16- (N=1159)
CD14- CD123- DR- CD16- CD4- CD8-
(76.0%)
Hu.CD8 | ADT_C0046
Hu.CD4_RPA.T4 | ADT_C0072
CD14- CD123- DR- CD16- CD4- CD8- (N=881
ILCs
(4.1%)
Hu.CD4_RPA.T4 | ADT_C0072
Hu.CD123 | ADT_C0064
CD19+ CD20+ (N=1174)
Naive B Cells
(70.2%)
Margina
(1.8%)
Memo
(7.8%
Hu.IgD | ADT_C0384
Hu.CD27 | ADT_C0154
Memory B Cells (N=92)
Plasmablasts
(8.7%)
IgD- Memory
(82.6%)
Hu.CD27 | ADT_C0154
Hu.CD20 | ADT_C0100
CD3+ TCRgd- (N=7237)
CD3+ CD56-
(87.3%)
CD3+ CD5
(12.4%)
Hu.CD8 | ADT_C0046
Hu.CD56 | ADT_C0047
CD4+ T Cells (N=3776)
Tregs(R3): 7.3%
Hu.CD127 | ADT_C0390
Hu.CD25 | ADT_C0085
CD8+ T Cells (N=1977)
R4(R4): 47.5%
R40(R40): 22.6%
R11(R11): 14.1%
Hu.CD45RA | ADT_C0063
Hu.CD62L | ADT_C0147
CD8+ CD62L- CD45RA- (N=447)
Early-like Effector CD8+(R8): 10.3%
Earl
Terminal Effector CD8+(R7): 19.5%
Inter
Hu.CD28 | ADT_C0386
Hu.CD27 | ADT_C0154
CD4+ CD62L- CD45RA- (N=747)
Early-like Effector CD4+(R48): 36.5%
Earl
Terminal Effector CD4+(R47): 3.9%
Inter
Hu.CD28 | ADT_C0386
Hu.CD27 | ADT_C0154
